# Deletion of the voltage-gated calcium channel, Ca_V_1.3, causes deficits in motor performance and associative learning

**DOI:** 10.1101/2020.08.12.248120

**Authors:** Marisol Lauffer, Hsiang Wen, Bryn Myers, Ashley Plumb, Krystal Parker, Aislinn Williams

**Affiliations:** Iowa Neuroscience Institute, University of Iowa, Iowa City, IA 52242, USA; Department of Psychiatry, University of Iowa, Iowa City, IA 52242, USA; Carver College of Medicine, University of Iowa, Iowa City, IA 52242, USA; Department of Physical Therapy and Rehabilitation Science, University of Iowa, Iowa City, IA 52242, USA

**Keywords:** calcium channels, motor learning, motor behavior, knockout mouse, Erasmus Ladder, rotarod, L-type channel, behavioral genetics, Cacna1d, learning

## Abstract

L-type voltage-gated calcium channels are important regulators of neuronal activity and are widely expressed throughout the brain. One of the major L-type voltage-gated calcium channel isoforms in the brain is Ca_V_1.3. Mice lacking Ca_V_1.3 are reported to have impairments in fear conditioning and depressive-like behaviors, which have been linked to Ca_V_1.3 function in the hippocampus and amygdala. Genetic variation in Ca_V_1.3 has been linked to a variety of psychiatric disorders, including autism and schizophrenia, which are associated with altered motor learning, associative learning, and social function. Here, we explored whether Ca_V_1.3 plays a role in these behaviors. We found that Ca_V_1.3 knockout mice have deficits in rotarod learning despite normal locomotor function. Deletion of Ca_V_1.3 is also associated with impaired gait adaptation and associative learning on the Erasmus Ladder. We did not observe any impairments in Ca_V_1.3 knockout mice on assays of anxiety-like, depression-like, or social preference behaviors. Our results suggest an important role for Ca_V_1.3 in neural circuits involved in motor learning and concur with previous data showing its involvement in associative learning.

## Introduction

Genome-wide association studies have identified many loci relevant to the risk for neuropsychiatric disorders, but the mechanisms by which these genes modify risk is unclear in most cases. One group of genes robustly linked to neuropsychiatric disease are the L-type voltage-gated calcium channel (LVGCC) genes, including *CACNA1D*. The *CACNA1D* gene encodes the pore-forming subunit of the L-type voltage-gated calcium channel, Ca_V_1.3, which is expressed in many tissues including the brain, heart, inner ear, and adrenal glands^1^. The *CACNA1D* gene has been linked to several neuropsychiatric disorders, including autism, bipolar disorder, depression, schizophrenia, and Parkinson’s disease^2-6^. Although the genetic connection between *CACNA1D* and neuropsychiatric disease is well-established, the functional role(s) of Ca_V_1.3 in different neural circuits remains under active investigation.

Loss of Ca_V_1.3 has been associated with multiple functional deficits in nervous system development and function. Due to its essential role in inner ear development^7^, Ca_V_1.3 knockout mice are congenitally deaf^8^, and therefore cannot undergo standard hearing-dependent associative learning paradigms, such as tone-paired fear conditioning^9^. In contrast, although Ca_V_1.3 influences light responsiveness in the retina^10,11^, mice lacking Ca_V_1.3 exhibit normal performance on vision-dependent tasks, such as the visible platform version of the Morris water maze^9,11^. Ca_V_1.3 is expressed in the hippocampus, and while Ca_V_1.3 KO mice have normal performance on the hidden platform version of the Morris water maze^9^ and the novel object recognition task^12^, they also have impaired object location memory^12,13^, all of which are hippocampus-dependent. Interestingly, there is evidence that object location memory also involves the cerebellum^14,15^. Some groups have identified abnormalities in anxiety- and depression-like behaviors in Ca_V_1.3 KO mice^11^, although not all groups have found similar deficits^9,13^. Pharmacological inhibition of Ca_V_1.3 in Ca_V_1.2 dihydropyridine insensitive mutant mice specifically in the ventral tegmental area caused no abnormalities in anxiety-like, depression-like, or social behaviors^16^.

Although *CACNA1D* is expressed in the striatum^17,18^ and cerebellum^19,20^, and has been associated with disorders that involve altered motor learning such as Parkinson disease and autism^5,21,22^, the specific role of Ca_V_1.3 in motor learning circuits has been relatively unexplored. Previous studies in Ca_V_1.3 KO mice have not shown deficits in locomotor function on fixed-speed rotarod^23^ or swim speed^9,11^, but one study using accelerating rotarod in a small sample showed a trend toward impaired motor learning over time^9^. We hypothesized that different behavioral tasks with larger sample sizes might reveal deficits in motor function and learning in Ca_V_1.3 KO mice. Here we explored the role of Ca_V_1.3 in motor, learning, and social behaviors, and found that Ca_V_1.3 KO mice have impairments in rotarod learning despite intact baseline locomotor function. We also find that Ca_V_1.3 KO mice display impaired associative learning and gait adaptation, a form of motor learning, on the Erasmus Ladder task without evidence of ataxia or motor incoordination. We find no deficits in Ca_V_1.3 KO mice in affective-like, anxiety-like, or social behaviors. Our results suggest that Ca_V_1.3 plays an important role in the neural circuits essential for motor and associative learning.

## Materials and Methods

### Animals

The generation of Ca_v_1.3 knockout (KO) mice (Cacna1d^tm1Jst^) has been described previously^8^. Breeding pairs of Ca_v_1.3^+/-^ (Hap) mice were maintained on a C57BL/6NTac background by crossing Hap offspring with C57BL/6NTac wild-type (WT) mice purchased from Taconic Biosciences (Rensselaer, NY). To generate experimental animals, Hap males were bred to Hap females to obtain male and female Ca_v_1.3 WT, Hap, and KO littermates. All mice were adults (at least 10 weeks old) at the time of testing. Behavioral experiments were run with two independent cohorts of mice in the following test order: cohort 1 (WT *n*=8 males and 7 females, Hap *n*=6 males and 9 females, KO *n*=6 males and 6 females) underwent Erasmus Ladder, rotarod, forced swim, and tail suspension; cohort 2 (WT *n*=8 males and 8 females, Hap *n*=8 males and 9 females, KO *n*=8 males and 7 females) underwent elevated zero maze, open field, 3-chamber social preference test, rotarod, tail suspension, and forced swim. All tests were performed with a minimum of two days between tests. Sample sizes are indicated in each figure. All experiments were carried out in a manner to minimize pain and discomfort, and animals were monitored after each experiment to ensure their health and safety. All experiments were conducted according to the National Institute of Health guidelines for animal care and were approved by the Institutional Animal Care and Use Committee at University of Iowa.

### Behavioral Procedures

General: Mice were same-sex group housed under regular light cycle lights on/off at 0900/2100 DST (0800/2000 non-DST). The average ambient temperature was 22°C and mice were provided with food and water *ad libitum*. All experiments were conducted during the animals’ light cycle. All equipment was cleaned between trials with 70% ethanol.

#### Open Field

Mice were placed in a 40 cm x 40 cm arena for 10 minutes under ∼115-130 lux. Activity was tracked by EthoVision software (Noldus, Leesburg, VA) and analyzed for total distance traveled and tendency to stay at the edge of the arena. For the latter, the arena was divided into the periphery and the center where each comprised 50% of the total surface area of the arena.

#### Rotarod

Mice were placed on the rotating drum of an accelerating rotarod (UGO Basile or IITC Life Science Mouse), and the time to fall or second passive rotation was recorded for each mouse. The speed of the rotarod accelerated from 4 to 40 rpm over a 5-minute period. Mice were given 3 trials/day for 5 days with a maximum time of 5 minutes, with at least a 10-minute inter-trial interval. Latency to fall or second passive rotation were recorded for each mouse each day.

#### Erasmus Ladder

Mice were trained on the Erasmus Ladder task (Noldus, Wageningen, The Netherlands) which has been described in detail elsewhere^24^. Briefly, the mice were trained on the Erasmus Ladder for 42 trials per day for a total of 4 consecutive days. Trials were separated by a random inter-trial interval ranging from 11-20 seconds. Data were analyzed for the following: number of trials where animal left on light cue, number of trials where animal left on air cue, number of trials where animal went onto the ladder before a cue was given, percentage of missteps, percentage of correct long steps (skipping at least one rung between steps), and percentage of correct short steps (using consecutive rungs).

#### Tail suspension test

Mice were suspended approximately 46-48 cm from the tabletop by lab tape wrapped around their tails, which was then attached to a hook on a horizontal rod to avoid bending the tail. Mice were video recorded for 6 minutes. Mice were removed from the apparatus if they climbed their tails to the top of the horizontal rod; data from these animals was excluded from analysis. Trials were analyzed for percentage of time immobile.

#### Forced swim test

Mice were placed in clear acrylic cylinders (outer diameter: 23 cm, inner diameter: 21.5 cm, height: 34 cm) filled halfway with water maintained at 20-25°C, and video recorded for 6 minutes. Trials were analyzed for latency to float and percentage of time immobile during the last 4 minutes of the trial. Mice that immediately floated upon placement into the water were not analyzed for latency but were analyzed for immobility.

#### Elevated zero maze

Mice were placed in a custom-built white plastic maze elevated 42.5 cm off the table with an internal diameter of 33.7 cm and outer diameter of 46 cm (internal pathway 5.8 cm wide). Walls on the closed sections were 10 cm high, and the lip on open sections was 0.6 cm high. Each mouse underwent a single 5-minute trial under ∼250 lux (open sections). Activity was tracked by EthoVision software for distance traveled, velocity, and duration spent in open/closed sections.

#### Three-chamber social preference test

Mice were placed in a matte, black plastic rectangular arena (L x W x H = 51 cm x 25.4 cm x 25.4 cm) divided into three compartments, with 10 cm wide opening between compartments and empty clear acrylic perforated cylinders in center of each outer compartment. Mice were habituated to the entire testing apparatus for 10 minutes. For the test, a novel conspecific mouse was placed under one cylinder while a novel object (colored plastic blocks) was placed under the other cylinder, and the test mouse was allowed to explore for 10 minutes. Mice were placed in middle compartment at the beginning of the test. Mice were removed from the apparatus if they climbed to the tops of the walls of the arena; data from these animals was excluded from analysis. Activity was tracked by EthoVision software for distance traveled, velocity, and interaction time (calculated by measuring the total duration during which the nose-point, but not the center-point, of the animal was within 1.5 cm of cylinder – a method designed by Benice and Raber^25^ to exclude instances where the mouse was rearing or climbing on the cylinder).

### Statistical analysis

All behavioral tasks except rotarod and Erasmus ladder were analyzed and graphed using GraphPad Prism 9.1 (GraphPad Software, San Diego, CA). Rotarod and Erasmus ladder data were analyzed via linear mixed effects modeling in RStudio with the lmerTest and emmeans packages and graphed in GraphPad Prism 9.1. Data are graphically represented as mean ± standard error of the mean (SEM) for each group. Data were analyzed with sex included as a factor to determine whether there were significant effects of sex. In cases where sex was not a significant factor, sexes were combined for analysis. Specific statistical tests used for each experiment are noted in the results section. Results were considered significant when p<0.05 (denoted in all graphs as follows: *p<0.05; **p<0.01).

## Results

### Mice lacking Ca_v_1.3 display normal locomotor and exploratory behavior

We first sought to determine whether Ca_v_1.3-deficient mice display abnormal locomotor and exploratory behaviors. In the open field task, we observed no differences in the distance traveled by Ca_v_1.3 Hap or KO mice compared to WT littermates (one-way ANOVA, F_2,45_=2.81, p=0.07) (Fig. 1A), nor a difference in the mean velocity of exploration (one-way ANOVA, F_2,45_=2.81, p=0.07) (Fig. 1B). We did observe a main effect of sex for distance traveled (two-way ANOVA, main effect of sex, F_1,42_=7.96, p=0.007) (Fig. S1A) and mean velocity (two-way ANOVA, main effect of sex, F_1,42_=7.96, p=0.007) (Fig. S1B); females traveled more and had higher velocities in general than males. When sex was included as a factor, there was no main effect of genotype for distance (two-way ANOVA, F_2,42_=3.08, p=0.057) or velocity (two-way ANOVA, F_2,42_=3.08, p=0.057) and no interaction effect (distance, two-way ANOVA, F_2,42_=0.57, p=0.57; velocity, two-way ANOVA, F_2,42_=0.57, p=0.57) (Fig. S1A-B). These data suggest that Ca_v_1.3 deficiency does not cause major impairments in baseline locomotion or exploration. We did not observe differences in animal weight at the beginning of behavioral testing (one-way ANOVA, F_2,87_=1.17, p=0.32) (Fig. S1D). The average age of mice at the beginning of behavioral testing did not differ between genotypes (one-way ANOVA, F_2,87_=0.15, p=0.86) (Fig. S1E).

**Fig. 1.**
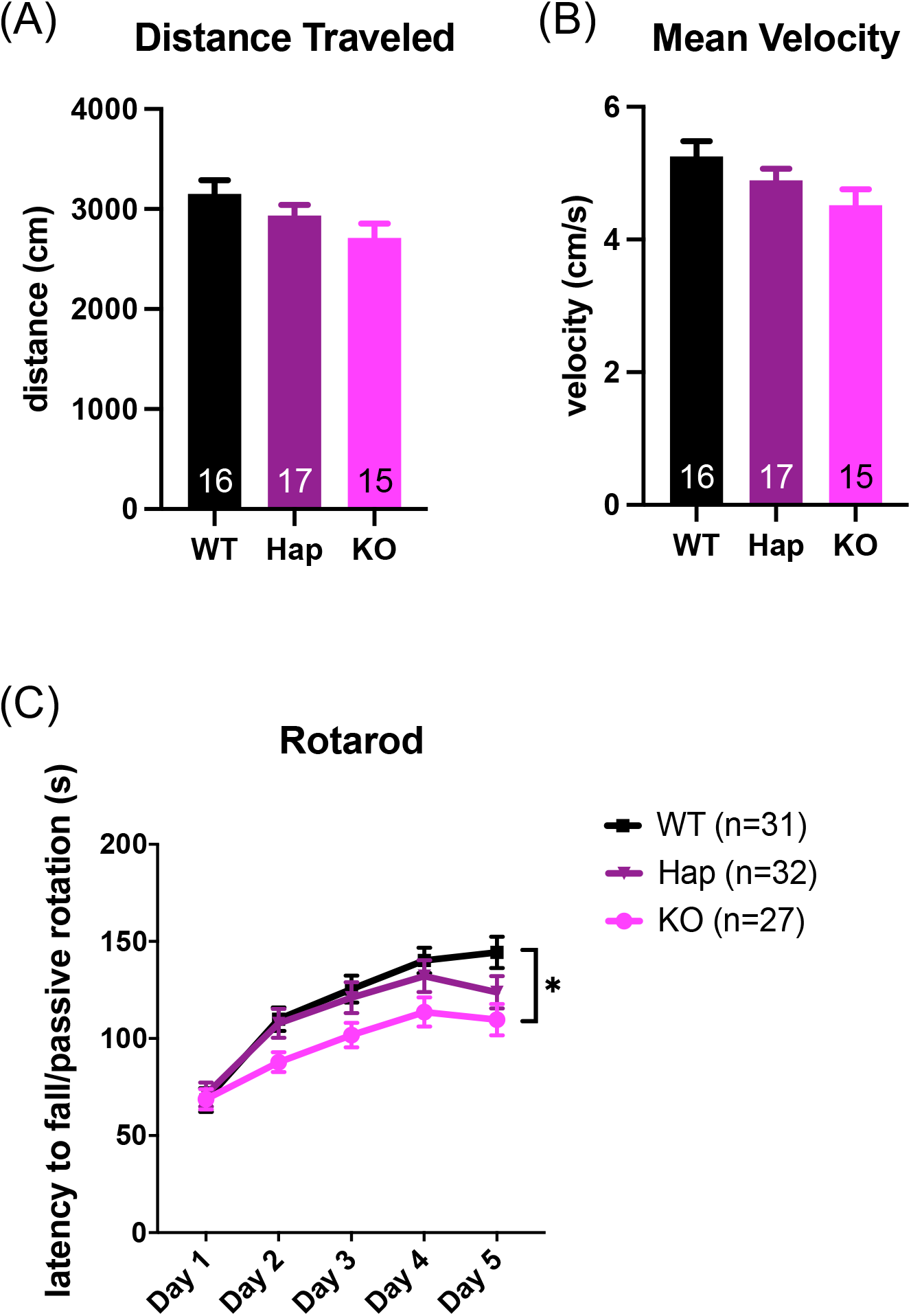
Ca_V_1.3 KO mice have normal locomotor exploration but impaired motor learning. **(A)** No differences were observed between genotypes in open field distance traveled. **(B)** No differences were observed between genotypes in mean velocity of movement in the open field. **(C)** KO mice perform significantly worse on the accelerating rotarod test of motor learning compared to WT mice. All genotypes improved over the course of training. A significant genotype x training interaction effect revealed that there was no significant effect of genotype on day 1, but WT was different from KO on days 2 through 5. The genotype effect on days 2 through 5 was always driven by KO mice performing significantly worse than WT mice (Hap mice were consistently indistinguishable from KO and WT mice on these days).

### Mice lacking Ca_v_1.3 display impaired motor performance

Although Ca_v_1.3 does not appear essential for basic locomotor and exploratory function (Figs. 1 and S1), we hypothesized that Ca_v_1.3 may be involved in motor learning, given its importance in other forms of learning. Therefore, we explored the ability of mice lacking Ca_v_1.3 to learn the accelerated rotarod task. We observed main effects of day of training (type III ANOVA with Satterthwaite’s method, main effect of training, F_4,564_=34.29, p<0.01) and sex (type III ANOVA with Satterthwaite’s method, main effect of sex, F_1,36_=4.54, p<0.05). We also observed a trend toward a genotype x day interaction effect (type III ANOVA with Satterthwaite’s method, main effect of sex, F_8,564_=1.73, p=0.08). Follow up testing showed that Ca_v_1.3 KO mice displayed impaired motor performance compared to their WT and Hap littermates on day 5 (Tukey’s posthoc test for genotype, p<0.01 for both WT v KO and Hap v KO on day 5) (Fig. 1C). This suggests that the general impairment shown by Ca_v_1.3 KO mice compared to WT mice is not likely due to ataxia or coordination issues as these would likely have caused KO mice to have impaired performance on day 1 (when no genotype effect was found). These data support the hypothesis that Ca_v_1.3 is relevant in motor learning.

Given the sex differences in the incidence of several disorders associated with Ca_v_1.3, such as autism and depression, we sought to determine whether sex differences were the primary drivers of the motor performance deficits we observed. When we analyzed rotarod data from male mice alone, we observed a genotype by day interaction effect (two-way repeated-measures ANOVA, F_8,164_=2.02, p=0.047) and a significant effect of training (two-way repeated-measures ANOVA, F_2.193,89.93_=43.55, p<0.01), but no main effect of genotype (two-way repeated-measures ANOVA, F_2,41_=2.51, p=0.09) (Fig. S1C). Post-hoc testing revealed that genotypes were not distinguishable on any day in males, although the differences between male WT and KO mice approached significance on day 4 (Tukey’s multiple comparisons test for genotype, p=0.05) and day 5 (Tukey’s multiple comparisons test for genotype, p=0.06). In female mice, we observed main effects of genotype (two-way repeated-measures ANOVA, F_2,43_=4.05, p=0.025) and day of training (two-way repeated-measures ANOVA, F_4,172_=42.0, p<0.01) but no interaction effect (two-way repeated-measures ANOVA, F_8,172_=0.73, p=0.66) (Fig. S1C). Post-hoc Tukey’s multiple comparisons test revealed a significant difference between female Hap and KO mice (p=0.03) and a trend toward a difference between WT and KO mice (p=0.08) (Fig. S1C). We conclude that the differences in rotarod performance are not likely to be driven primarily by sex differences.

### Ca_v_1.3 knockout mice display associative learning deficits

The Erasmus Ladder task permits assessment of a variety of behaviors, including associative learning (ability to learn visual and sensory start cues), gait adaptation (learning to use longer steps to cross the ladder), cerebellum-dependent associative motor learning (learning to time a jump for an auditory cue), and motor coordination (missteps). The task parameters have been described in detail elsewhere^24,26^. Briefly, in this task, mice start a motor trial after a cue (a bright light, followed by air if they do not begin within 3 seconds of the light coming on). The apparatus measures the animal’s steps across a discontinuous ladder and detects errors for each trial (there are 42 trials in a session). We could not assess cerebellum-dependent tone-cued associative learning with this task because the conditioned stimulus is a tone and Ca_v_1.3 KO mice are congenitally deaf. However, since Ca_v_1.3 KO mice have normal vision and mechanosensation^11,23^, we were able to assess light- and air-cued associative learning, motor coordination, and gait adaptation while mice learned to cross the ladder. Most animals tended not to start the task following the light cue with no differences between genotypes (type III ANOVA with Satterthwaite’s method, main effect of genotype, F_2,36_=1.37, p=0.27) (Fig. 2A). However, WT and Hap mice generally started the task following the air cue, while Ca_V_1.3 KO mice responded to this cue less frequently (type III ANOVA with Satterthwaite’s method, main effect of genotype, F_2,36_=58.36, p<0.01; both WT and Hap mice differed significantly from KO by Tukey’s posthoc multiple comparisons test p<0.01; no difference between WT and Hap mice) (Fig. 2B). Ca_v_1.3 KO mice also walked onto the ladder before any cue was given (i.e., they did not wait for the cue to start the trial) more often than their WT and Hap littermates (type III ANOVA with Satterthwaite’s method, main effect of genotype, F_2,39_=55.57, p<0.01; genotype by session interaction, F_6,108_=2.18, p<0.05). Follow up testing for the genotype by session interaction effect showed that WT and Hap were different from KO for each session (p<0.01) with no difference between WT and Hap mice during any session (Fig. 2C). These data support previous studies showing that Ca_v_1.3 is important for associative learning.

**Fig. 2.**
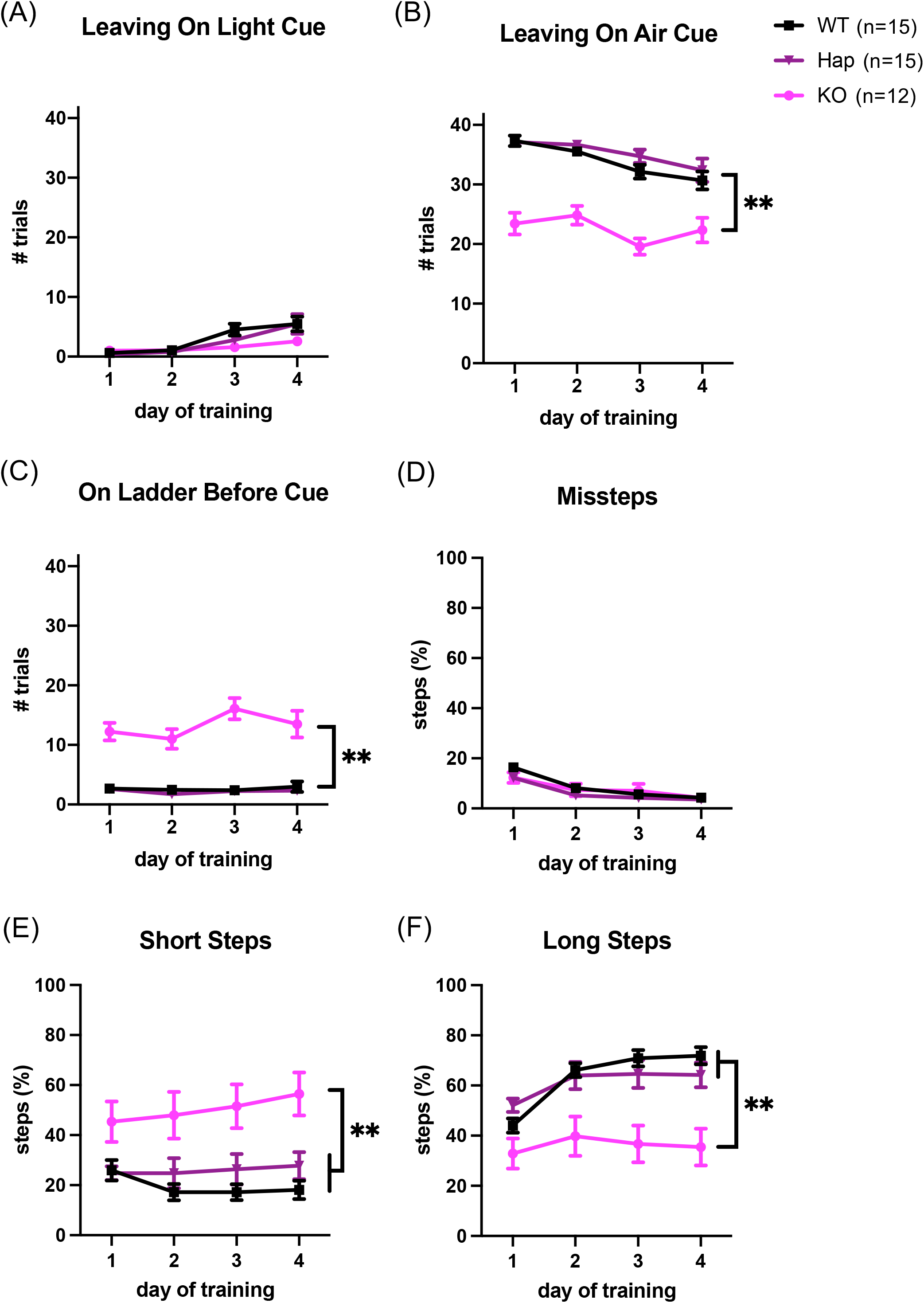
Ca_V_1.3 KO mice display impaired associative learning and gait adaptation on the Erasmus Ladder task. **(A)** All mice demonstrate low propensity to leave the start box on the light cue; no differences noted between genotypes. **(B)** KO mice are significantly less likely to leave the start box with the air cue compared to WT and Hap littermates. **(C)** KO mice are significantly more likely to move onto the ladder prior to any start cue compared to WT and Hap littermates. **(D)** All animals have a reduction in missteps over time. There is a significant genotype by day interaction effect but Tukey’s multiple comparisons test identified no differences between genotypes on any individual day. **(E)** WT and Hap mice use short steps to cross the ladder less frequently than KO mice. There is also a genotype x day interaction effect; follow up testing shows that WT do not differ from KO on day 1 but do on days 2 through 4, suggesting that WT mice learn to use fewer short steps over time, but KO mice do not. **(F)** WT and Hap mice differ in their use of long steps compared to KO mice. There is also a genotype x day interaction effect; follow up testing shows that WT do not differ from KO on day 1 but do on days 2 through 4, suggesting that WT mice learn to use long steps over time, but KO mice do not.

### Ca_v_1.3 knockout mice display deficits in gait adaptation

Interestingly, once on the ladder, Ca_V_1.3 KO mice did not display an increase in missteps compared to WT or Hap mice, suggesting that loss of Ca_v_1.3 does not cause ataxia or coordination problems (type III ANOVA with Satterthwaite’s method, main effect of day, F_3,108_=94.42, p<0.01, main effect of genotype, F_2,36_=0.79, p=0.46) (Fig. 2D). There is a significant genotype by day interaction effect (type III ANOVA with Satterthwaite’s method, genotype by day interaction effect, F_6,108_=2.88, p<0.05) but Tukey’s multiple comparisons test identified no differences between genotypes on any individual day. The normal motor coordination of Ca_V_1.3 KO mice on the motor aspects of the Erasmus Ladder further supports the notion that the observed rotarod motor deficits are not secondary to incoordination.

In agreement with the rotarod data, Ca_V_1.3 KO mice displayed impairments in gait adaptation, which is a form of motor learning. Typically, as mice learn the Erasmus Ladder task over several days, they transition from using short steps to using longer steps to cross^27,28^, since long steps are a more efficient strategy. For short steps, we observed a main effect of sex (type III ANOVA with Satterthwaite’s method, main effect of sex, F_1,36_=12.47, p<0.01) and a genotype by sex interaction (type III ANOVA with Satterthwaite’s method, genotype by sex interaction, F_2,36_=3.60, p<0.05). Follow up testing showed that there were no differences in short steps in male mice by genotype (WT v KO, p=0.30, Hap v KO, p=0.48) (Fig. S2A), but female KO mice differed from both female WT and Hap mice (p<0.01) (Fig. S2B). We also observed a main effect of genotype (type III ANOVA with Satterthwaite’s method, main effect of genotype, F_2,36_=11.94, p<0.01) and a genotype by day interaction effect (type III ANOVA with Satterthwaite’s method, F_6,108_=2.36, p<0.01) (Fig. 2E); post-hoc testing showed that WT mice and Hap mice were different from KO mice on all days of testing (p<0.05 for day 1 and p<0.01 for days 2 through 4). For long steps, we observed main effects of genotype (type III ANOVA with Satterthwaite’s method, main effect of genotype, F_2,36_=13.79, p<0.01), sex (F_1,36_=11.57, p<0.01) and day of training (F_3,108_=24.69, p<0.01), as well as a genotype by day interaction effect (F_6,108_=5.22, p<0.01) (Fig. 2F). In contrast to WT mice, Ca_V_1.3 KO mice did not increase their use of long steps over consecutive days (Tukey’s multiple comparisons test, day 1 p=0.21, days 2 through 4 p<0.01) (Fig. 2F). Overall, female mice were less likely to use long steps than male mice (p<0.01). Taken together, these results show that Ca_V_1.3 KO mice have an impairment in motor learning on the Erasmus ladder task.

### Mice lacking Ca_v_1.3 do not display depression-like or anxiety-like behaviors

Since genetic variation in Ca_V_1.3 has been associated with mood disorders in humans^3,4,6^ and antidepressant-like behavior in male mice^11^, we explored whether mice lacking Ca_V_1.3 have abnormal depression-like or anxiety-like behaviors. In the tail suspension test, which has been used as a predictor of antidepressant efficacy^29^, we observed no differences in time spent immobile between genotypes (one-way ANOVA, F_2,85_=0.33, p=0.72) (Fig. 3A). When sexes were separated, no differences between sexes or genotypes were observed (two-way ANOVA, main effect of genotype F_2,82_=0.34, p=0.71; main effect of sex, F_1,82_=0.001, p=0.97; interaction effect F_2,82_=1.36, p=0.26) (Fig. S3A). Similarly, in the forced swim test of behavioral despair^29,30^, we found no differences between genotypes in percent time spent immobile (one-way ANOVA, F_2,87_=1.81, p=0.17) (Fig. 3B) or in latency to begin floating (one-way ANOVA, F_2,81_=0.61, p=0.55) (Fig. 3C). When sex was included as a factor, we observed no differences between genotypes for immobility (two-way ANOVA, main effect of genotype F_2,84_=1.73, p=0.18; main effect of sex, F_1,84_=0.05, p=0.83; interaction effect F_2,82_=0.63, p=0.54) (Fig. S3B) or latency to float (two-way ANOVA, main effect of genotype F_2,78_=0.83, p=0.44; main effect of sex, F_1,78_=2.21, p=0.14; interaction effect F_2,78_=0.82, p=0.44) (Fig. S3C).

**Fig. 3.**
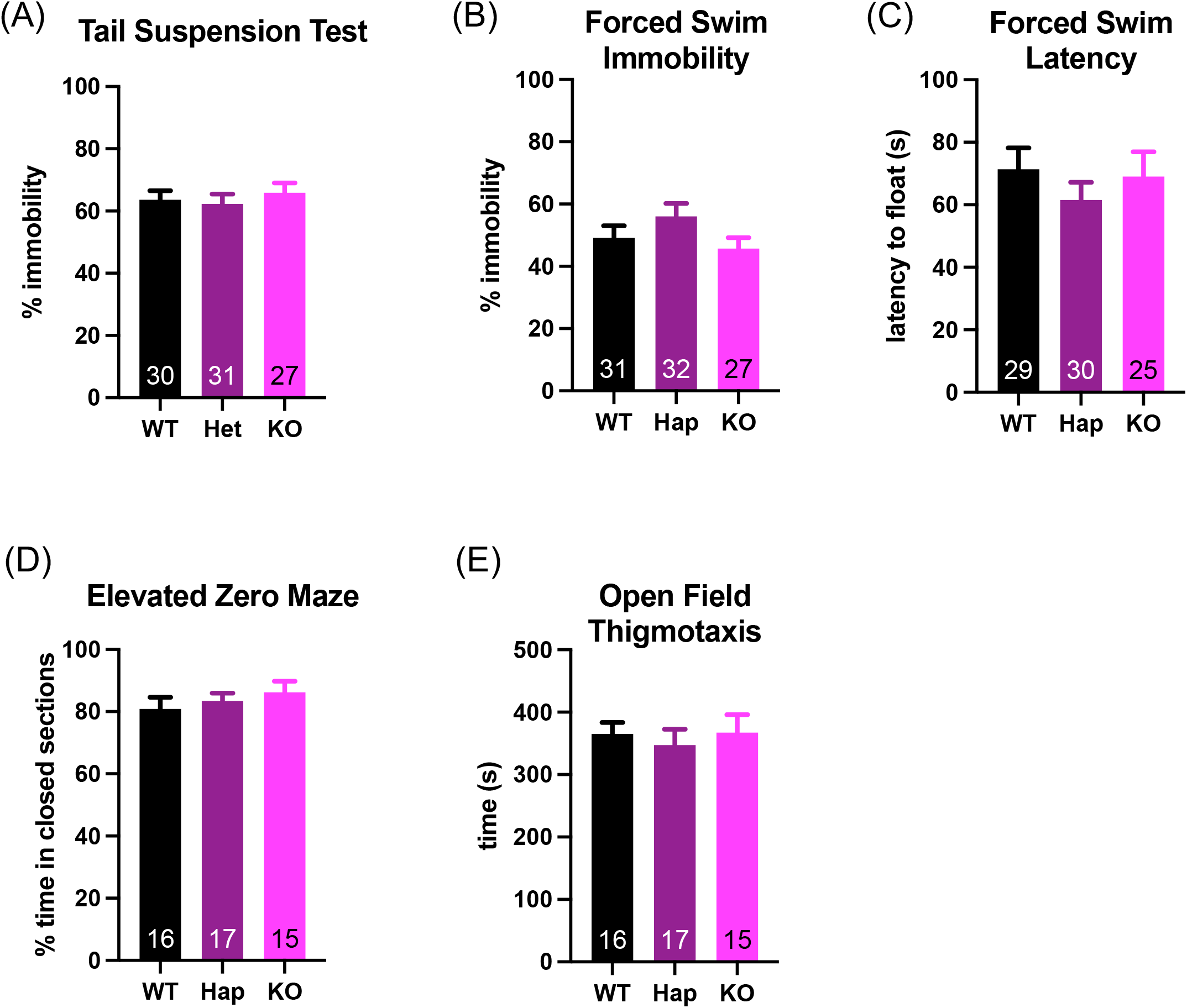
Ca_V_1.3 KO mice have no deficits in affective or anxiety-like behaviors. **(A)** KO mice are not different from WT and Hap littermates in time spent immobile on the tail suspension test. KO mice do not differ from WT and Hap littermates in the forced swim test in either (**B**) percent time spent immobile or (**C**) latency to start floating. KO mice do not differ from WT and Hap littermates in anxiety-like behaviors in either (**D**) the elevated zero maze (measured as time spent in closed segments) or (**E**) the open field test (measured as time spent in the periphery of the arena).

We examined anxiety-like behaviors using the elevated zero maze and the open field task, both of which measure the animal’s propensity to explore a riskier part of the environment. In the elevated zero maze, we found no differences in percent time spent in the closed segments by genotype (one-way ANOVA, F_2,45_=0.65, p=0.53) (Fig. 3D); the same was true when sex was included as a factor (two-way ANOVA, main effect of genotype, F_2,42_=0.39, p=0.53) (Fig. S3D). We observed no differences between genotypes in distance traveled (one-way ANOVA, F_2,45_=2.50, p=0.09) (Fig. S4A) or mean velocity (one-way ANOVA, F_2,45_=2.47, p=0.10) (Fig. S4B) in the elevated zero maze. When sex was included as a factor in the elevated zero maze, we still observed no differences in distance traveled (two-way ANOVA, main effect of genotype F_2,42_=2.36, p=0.11; main effect of sex, F_1,42_=0.44, p=0.51; interaction effect F_2,42_=0.70, p=0.50) (Fig. S4C) or velocity (two-way ANOVA, main effect of genotype F_2,42_=2.34, p=0.11; main effect of sex, F_1,42_=0.43, p=0.52; interaction effect F_2,42_=0.70, p=0.50) (Fig. S4D). In the open field task, time spent in the center of the field is associated with reduced anxiety, whereas time spent in the periphery (also called thigmotaxis) is associated with higher anxiety^31,32^. In the open field test, we observed no differences in thigmotaxis with sexes combined (one-way ANOVA, main effect of genotype, F_2,45_=0.21, p=0.81) (Fig. 3E) or with sex included as a factor (two-way ANOVA, main effect of genotype, F_2,42_=0.23, p=0.79) (Fig. S3E).

### Ca_v_1.3 knockout mice display normal preference for social interaction

Since rare mutations in Ca_v_1.3 have been linked to autism, we wanted to test whether loss of Ca_v_1.3 would impair social preference. We tested this using the three-chamber social preference test^33,34^. In the social preference test, we found that mice lacking one or both copies of Ca_v_1.3 prefer to explore a cylinder containing another mouse rather than one containing an object, as do their WT and Hap littermates (two-way ANOVA, main effect of test object, F_1,86_=47.19, p<0.01) (Fig. 4A). Ca_v_1.3 Hap mice explored both cylinders more, on average, than WT and KO mice, while WT and KO mice were not different from each other (two-way ANOVA, main effect of genotype, F_2,86_=5.80, p<0.01, Tukey’s multiple comparison test WT vs Hap and Hap vs KO p<0.05). There was no interaction effect (two-way ANOVA, F_2,86_=1.56, p=0.22). We found no genotype-specific differences in time spent exploring the arena as a whole during the task (one-way ANOVA, main effect of genotype, F_2,44_=0.46, p=0.63) (Fig. 4B) nor in movement velocity during exploration (one-way ANOVA, main effect of genotype F_2,44_=0.47, p=0.63) (Fig. 4C), supporting the open field and elevated zero maze data showing no differences in basal locomotor function or velocity (Fig. 1A-B and S4).

**Fig. 4.**
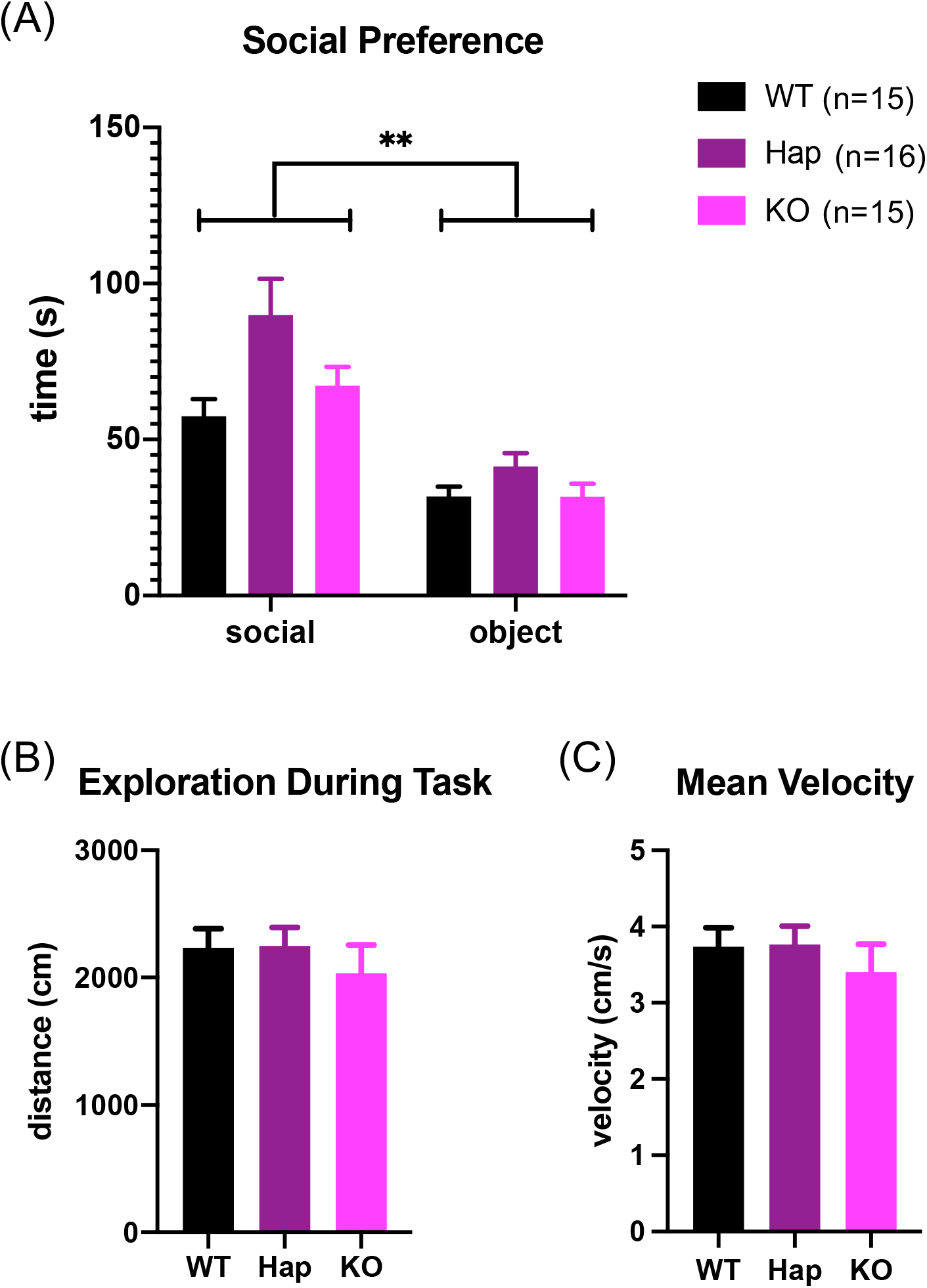
Ca_V_1.3 KO mice do not display social interaction deficits. **(A)** All groups display a preference for interaction with a novel mouse over a novel object. Hap mice spend more time interacting with the test objects overall than their WT and KO littermates. There was no interaction effect. There are no differences between genotypes in (**B**) total exploration (arena plus objects) during the task or (**C**) velocity of movements during the task.

## Discussion

L-type voltage-gated calcium channels (LVGCCs) have important roles in learning and memory and have been linked to multiple human neuropsychiatric diseases. Here we investigated the behavioral phenotypes caused by loss of one specific LVGCC, Ca_V_1.3. We found that complete loss of Ca_V_1.3 is associated with deficits in the accelerating rotarod task (Fig. 1C), indicative of impaired motor learning or performance. These deficits were not present on day 1, as may have been expected if they were due to ataxia and incoordination. We also observed deficits in a second form of motor learning, gait adaptation on the Erasmus Ladder, in Ca_V_1.3 KO mice (Fig. 2E-F). We identified abnormalities in light- and air-cued associative learning on the Erasmus Ladder task in Ca_V_1.3 KO mice (Fig. 2), which concurs with previous studies demonstrating the importance of Ca_V_1.3 in associative learning. These data taken together with normal locomotor exploratory behavior (Figs. 1, 4, S1, and S4) and lack of ataxia or incoordination on Erasmus Ladder (Fig. 2D) suggest that the rotarod and gait adaptation deficits observed in Ca_V_1.3 KO mice are not related to underlying deficits in locomotor ability, but rather, demonstrate a deficit in motor performance or learning. Several other mouse lines with Purkinje cell dysfunction display the same gait adaptation phenotype, i.e. they persistently use short steps rather than switching to using long steps on the Erasmus Ladder^27,35^. These data, in combination with the recent report that L-type calcium channels are involved in GABA release from molecular layer interneurons onto Purkinje cells^36^, raise the possibility that Ca_V_1.3 functions in cerebellar cortical circuits that regulate Purkinje cell activity.

Our data differ from a previous study of Ca_V_1.3 KO mice which reported no statistical difference in accelerating rotarod performance in Ca_V_1.3 KO mice on a C57BL/6:129Sve F2 hybrid background ^9^. We used mice on a pure C57BL6/Ntac genetic background and significantly larger sample sizes, which may account for the difference in results. The only other published rotarod experiment in Ca_V_1.3 KO mice used the same genetic background as our animals (C57BL6/NTac) in a single trial of the rotarod set at a fixed speed of 18rpm^23^, which showed no difference between wild-type and Ca_V_1.3 KO mice. When used in a single trial at a fixed speed, the rotarod task assesses motor coordination but not motor learning over time^37-39^; thus our rotarod data measure a different behavioral outcome than the study by Clark and colleagues. Interestingly, our data show that Ca_V_1.3 KO mice are indistinguishable from WT mice on day 1 of rotarod testing, and differences only emerged at later days of testing. Our results regarding normal locomotor activity and exploration agree with those of earlier studies^11,23^.

We did not observe depression-like behaviors in Ca_V_1.3 KO mice as has been previously described^11^. Notably, other studies of this animal model have also not detected depression-like or anxiety-like behaviors^9,13^. The origin of this discrepancy is unclear, and merits further investigation. As mentioned above, we used larger sample sizes as compared to these previous studies, which may partially explain some of the differences between our results and those of other groups.

Our Erasmus Ladder data support the hypothesis that Ca_V_1.3 is important for associative learning when mildly aversive stimuli (bright light and air puffs) are used. These data are consistent with the findings of McKinney and Murphy^9^ where Ca_V_1.3 deletion disrupted consolidation but not extinction of contextual fear conditioning when a mild foot shock stimulus was used for training (single 2-sec 0.50 mA shock). Notably, when Ca_V_1.3 KO mice are exposed to stronger fear conditioning stimuli (five footshocks of 0.7 mA) there appear to be no differences in either consolidation or extinction of fear learning^40^, suggesting that Ca_V_1.3 is not essential for all forms of aversive learning but may modulate learning under less stressful circumstances. Conversely, one study suggests that activating Ca_V_1.3 specifically in the ventral tegmental area contributes to cocaine preference while also inducing depressive-like behavior and social preference deficits^16^. There is now also a mouse line which overexpresses Ca_V_1.3 in excitatory forebrain neurons^41^ which might reveal whether Ca_V_1.3 in specific brain regions or neuronal subtypes is associated with specific types of behavioral circuit dysfunction. For example, it would be interesting to study whether Ca_V_1.3 plays a role in milder forms of reward learning, such as operant conditioning paradigms, as different forms of learning may be mediated by different neural circuits.

Notably, since our approach utilizes global knockout mice where Ca_V_1.3 is deleted from birth, we cannot rule out the possibility that unidentified compensatory changes may mask other roles for Ca_V_1.3 in the brain. In other words, the absence of a phenotype in these assays does not imply that Ca_V_1.3 is not involved in the specified behaviors, since developmental compensatory mechanisms may support those functions when Ca_V_1.3 is absent (such as upregulation of other calcium channels^42,43^). Recent work shows that L-type calcium channels can regulate GABAergic signaling between molecular layer interneurons and Purkinje cells in the cerebellar cortex^36^, and L-type channels have also been implicated in GABA and dopamine release in the striatum^44-47^. Both Ca_V_1.2 and Ca_V_1.3 are expressed in the striatum^17,18^ and the cerebellum^19,20^ but currently available pharmacologic agents cannot distinguish between these channels, therefore careful genetic work will be necessary to dissect their respective roles in specific brain regions and cell types. Future work should focus on acute disruption of Ca_V_1.3 expression using cell-type or region-specific methods to determine whether additional Ca_V_1.3-dependent phenotypes can be elicited.

In summary, our data demonstrate the importance of Ca_V_1.3 in both motor and associative learning. Abnormalities in motor and associative learning have been demonstrated in multiple neuropsychiatric disorders, including autism, bipolar disorder, and schizophrenia. Interestingly, despite the association of Ca_V_1.3 with mood disorders in humans, we do not find evidence of affective or anxiety-like abnormalities in mice lacking Ca_V_1.3. This suggests that perhaps Ca_V_1.3 contributes to common cognitive deficits across neuropsychiatric disorders. Therefore, therapies that aim to modulate Ca_V_1.3 function may have broad applicability across neuropsychiatric conditions.

## Acknowledgements

The authors would like to thank Dr. Shane Heiney and the Neural Circuits and Behavior Core at the University of Iowa, Dr. Kumar Narayanan, and Parker Abbott for training and behavioral equipment used in this study. We also thank Dr. Geoffrey Murphy for Ca_V_1.3 knockout mice, and the members of the Iowa Neuroscience Institute for thoughtful comments on this manuscript. This project was funded by NIH R01MH118240 (K.L.P.), K01MH106824 (K.L.P.), KL2TR002536 (A.J.W.), the Roy J. Carver Charitable Trust (A.J.W.), the Graduate Program in Neuroscience (A.P.), and the Roy J. and Lucille A. Carver College of Medicine (B.M.). The authors declare that they have no conflicts of interest.

## Data Sharing/Availability Statement

The data that support the findings of this study are available from the corresponding author upon reasonable request.

## Figure Legends

**Fig. S1** Data from Figure 1 separated by sex. Overall, female mice cover statistically more distance (**A**) at a greater speed (**B**) than male mice in the open field task. There is no main effect of genotype for either distance or velocity and there is no interaction effect. **(C)** On the rotarod task for male mice, there is a main effect of day of training and a genotype by day interaction effect but no main effect of genotype. In males, Tukey’s multiple comparisons test found no significant differences between genotypes on any individual day. For females on the rotarod task, there was a main effect of genotype; follow up testing with Tukey’s multiple comparisons test identified a significant difference between Hap and KO female mice. **(D)** There were no differences between genotypes for animal weight. **(E)** The mean age of animals at the beginning of behavioral experiments did not differ between genotypes.

**Fig. S2** Data from Figure 2 separated by sex. **(A)** There are no statistically significant genotype effects for males in the use of short steps on the Erasmus Ladder. **(B)** Female KO mice use short steps more often on the Erasmus Ladder compared to their WT and Hap littermates.

**Fig. S3** Data from Figure 3 separated by sex. There are no sex or genotype effects for any affective or anxiety-like task. **(A)** There are no sex or genotype effects for time spent immobile in the tail suspension test. **(B)** There are no sex or genotype effects for percent time spent immobile in the forced swim test. **(C)** There are no sex or genotype effects for latency to float in the forced swim test. **(D)** There are no sex or genotype effects for time spent in closed segments of the elevated zero maze. **(E)** There are no sex or genotype effects in the proportion of time spent in the outer edge of the arena in the open field test.

**Fig. S4** Additional data from the elevated zero maze. KO mice are not different from WT and Hap littermates in terms of (**A**) distance traveled or (**B**) velocity of movement on the elevated zero maze. There is no main effect of sex on elevated zero maze in (**C**) distance traveled or (**D**) velocity.

