## Supplementary figures and images for "Deletion of the voltage-gated calcium channel, Ca_V_1.3, causes deficits in motor performance and associative learning"

### Supplemental Figure 1

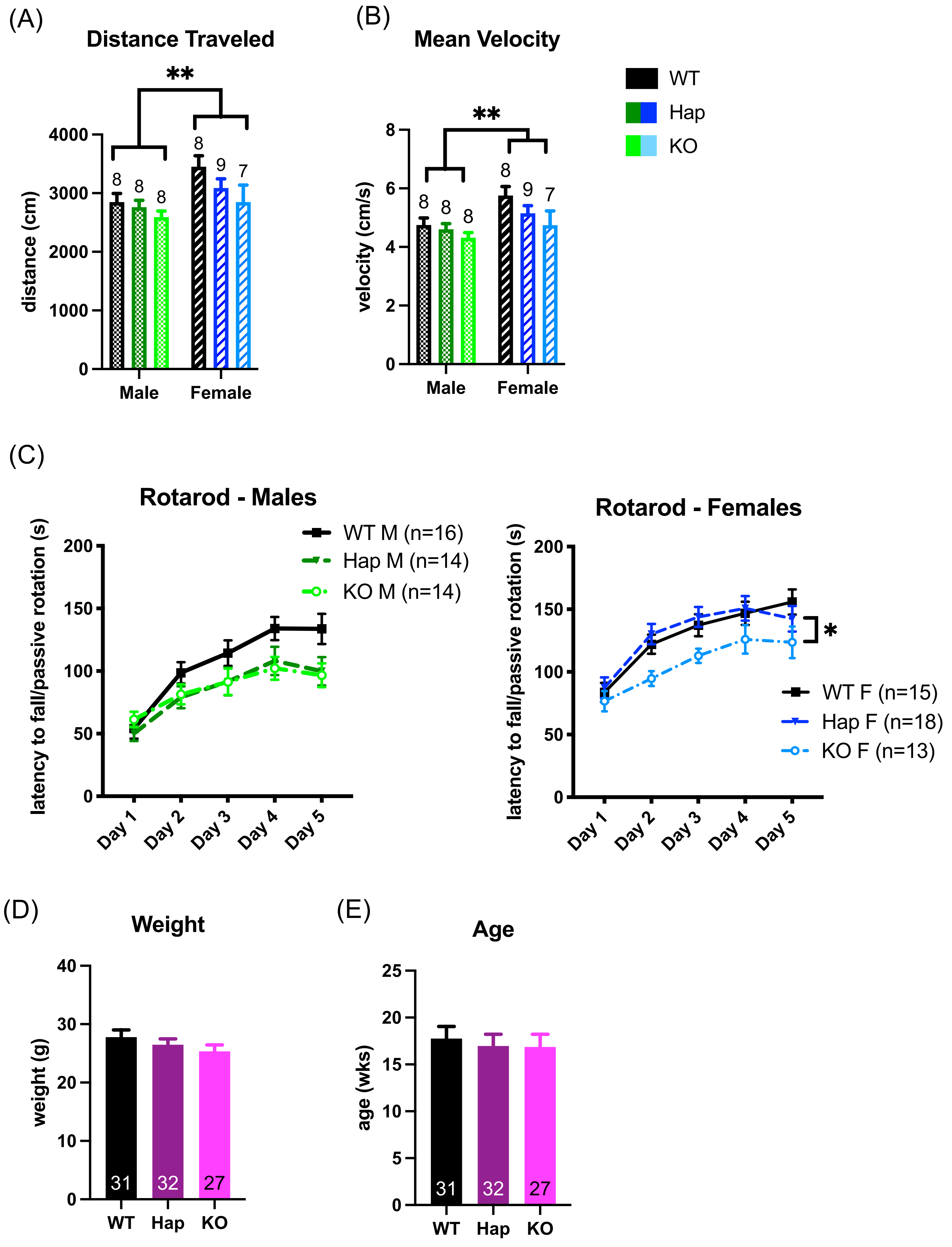

### Supplemental Figure 2

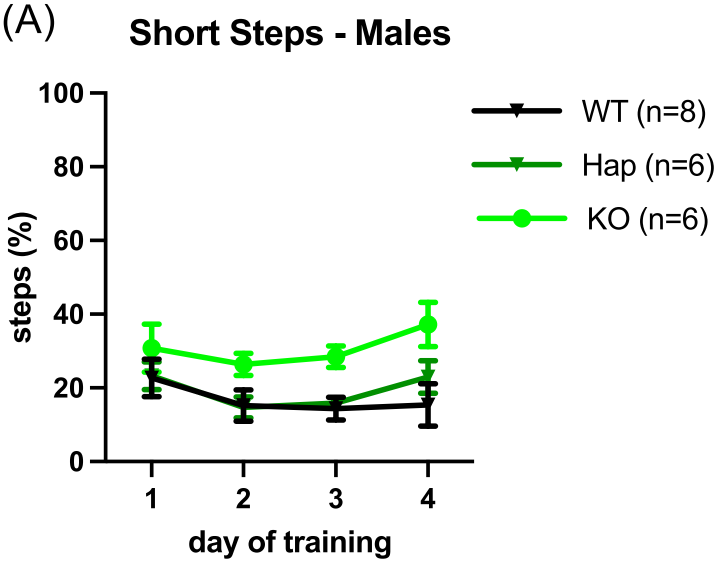

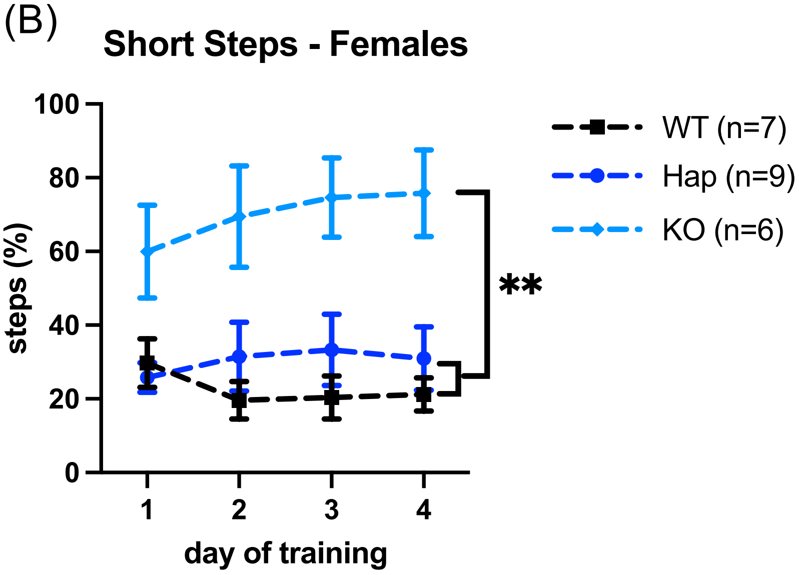

### Supplemental Figure 3

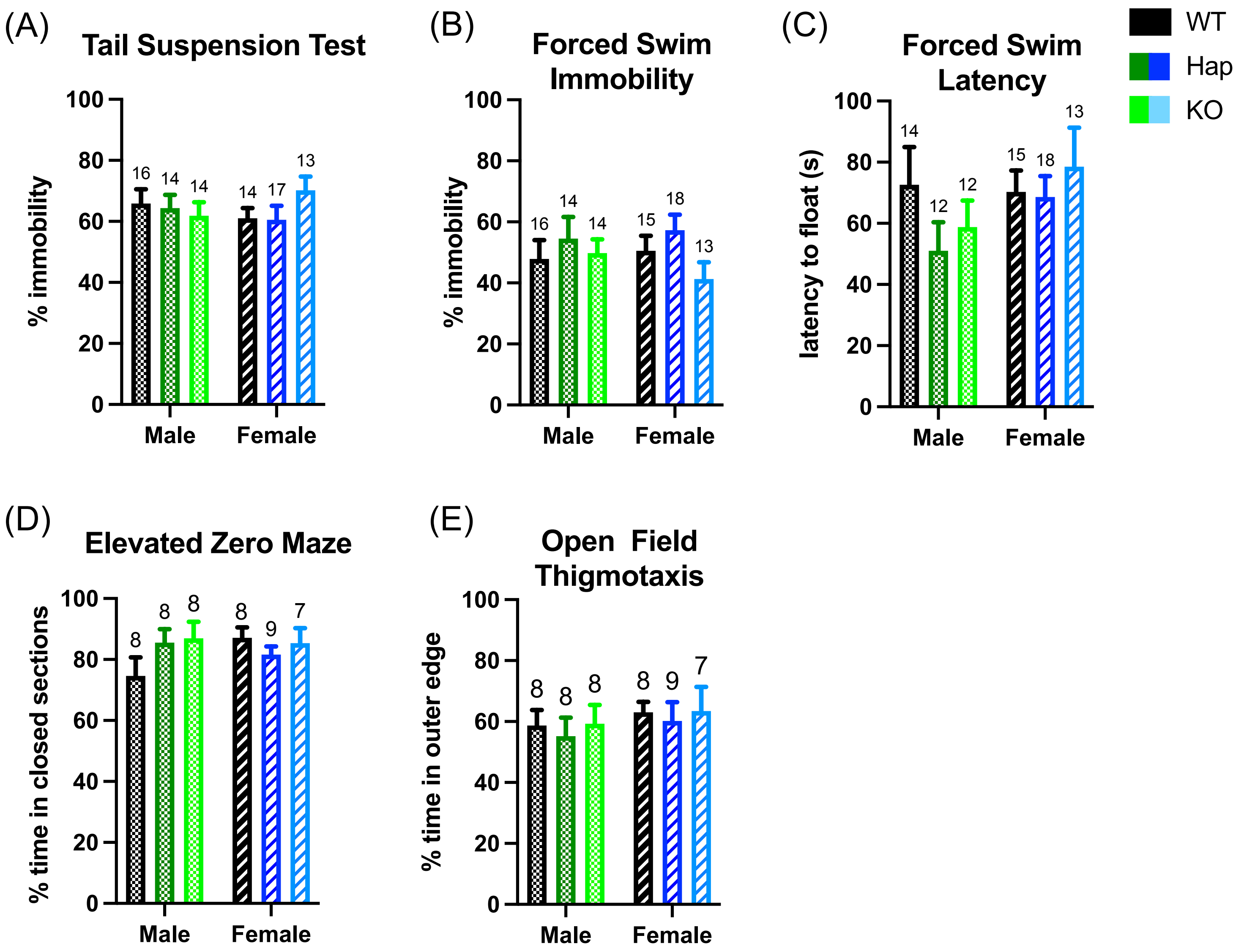

### Supplemental Figure 4

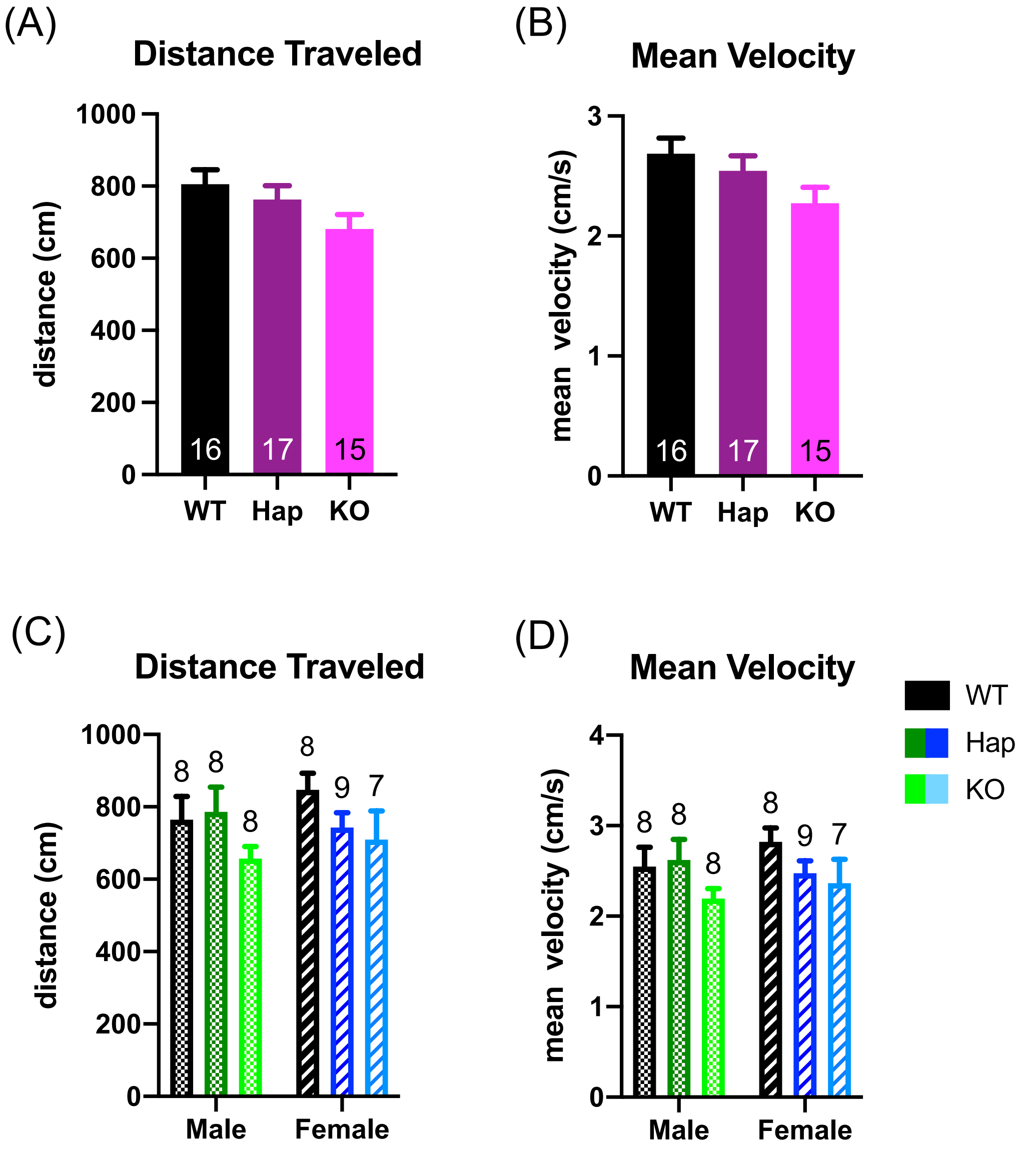
